# Validation of deep learning enabled software MetronMind to measure vertebral heart size and vertebral left atrial size in dogs

**DOI:** 10.64898/2025.12.07.692858

**Authors:** K. Tess Sykes, Sonya G. Gordon, John J. Craig, Sonya Wesselowski, Alice Watson

## Abstract

Vertebral heart size (VHS) and vertebral left atrial size (VLAS) are objective radiographic measurements of heart and left atrial size respectively and are associated with inter and intraobserver variability when measured by humans. Artificial intelligence (AI) tools to determine VHS and VLAS have been developed which may reduce variability and save time. Two manual methods for measuring VHS and VLAS on right and left lateral canine thoracic radiographs were compared. Measurements of VHS and VLAS made by deep learning enabled program, MetronMind, were compared to a trained observer on right and left lateral radiographs from 1058 client-owned dogs including 80 breeds with a variety of heart sizes, thoracic conformations and radiographic quality. This was a retrospective, single center, method comparison study. Pearson’s correlation, Bland-Altman plots and Passing-Bablok regression were used to assess agreement.

Correlation between traditional and modified manual measurements for VHS and VLAS were strong (r=0.994 and r=0.974 respectively), with minimal bias (−0.10 and 0.04 vertebrae respectively) indicating that the modified methods closely approximate traditional measurements obtained from right lateral views. MetronMind measurements of VHS and VLAS from right lateral radiographs correlated well with the human observer’s modified measurements (r=0.947 and r=0.811 respectively), showing small mean biases (0.08 and 0.07 vertebrae respectively). Correlation between left and right lateral radiographic measurements of VHS (0.87 and 0.91) was higher than for VLAS (0.73 and 0.64) and bias was larger for VHS (0.26 and 0.31 vertebrae) than VLAS (-0.13 and -0.10 vertebrae) for humans and MetronMind respectively. MetronMind can therefore assist veterinarians with measuring VHS and VLAS in dogs and right lateral radiographs are preferred. Future studies are needed to compare artificial intelligence derived radiographic measures with echocardiographic measures of cardiac size.

## Introduction

Clinicians consider transthoracic echocardiography the clinical gold standard for assessing heart size in dogs [1]. Where echocardiograms are unavailable, thoracic radiographs provide a reasonable substitute for diagnosing, staging and monitoring cardiac disease [2, 3]. Assessment of heart size on thoracic radiographs was historically based on how many intercostal spaces the heart spans within the thorax on a lateral radiograph, how much of the thoracic space the heart occupies or other subjective impressions [4]. Variability in thoracic confirmation in dogs, coupled with phase of respiration or oblique positioning during exposure results in inaccuracies and inconsistent interpretations of heart size [5].

Vertebral heart size (VHS) provides a quantitative measurement of heart size typically performed on right lateral thoracic radiographs [5]. The VHS method adopts the principle that a fixed relationship exists between heart size, and the length of the thoracic vertebral bodies in various breeds. Furthermore, it provides a consistent reference measurement (the vertebrae) allowing clinicians to compare cardiac measurements over time. Obtaining an accurate traditional VHS measurement requires identification of five anatomic landmarks within the thorax: the central and ventral aspect of the carina, the ventral most aspect of the cardiac apex, the caudal vena cava, the cranial cardiac waist, and the cranial aspect of the 4^th^ thoracic vertebra [5, 6]. The major axis or height of the heart is drawn from the central and ventral aspect of the carina to the left apex. The minor axis or width of the heart drawn roughly perpendicular to the height line at the widest aspect. The width can be standardized by orientation to the caudal vena cava, typically mid or ventral boarder. Two lines of the same length as the height and width are drawn parallel to the vertebral bodies, beginning with the cranial edge of the 4^th^ thoracic vertebra. A numerical VHS value is calculated by adding measurements of the height and width of the cardiac silhouette and converted to vertebral body units to the nearest 0.1 vertebrae [5]. Measuring VHS is useful for identifying cardiomegaly in dogs [7] and discriminating between stage B1 and B2 myxomatous mitral valve disease where echocardiography is not feasible [8], and sequential measurements can help predict the onset of congestive heart failure [9, 10]. As with subjective assessments, the accuracy of VHS measurements depends on the quality of the radiographic technique, patient positioning and clinician experience [6], and is subject to intraobserver and interobserver variability [6].

Vertebral left atrial size (VLAS) is a radiographic surrogate for measurement of left atrial size. Clinicians typically measure VLAS on a right lateral view of the thorax by drawing a line from the center of the most ventral aspect of the carina to the most caudal aspect of the left atrium where it intersects with the dorsal border of the caudal vena cava. Like VHS, a line of the same length is drawn parallel to the vertebral bodies, beginning with the cranial edge of the 4^th^ thoracic vertebra [11]. The smaller absolute measurement of VLAS results in larger intraobserver and interobserver variability than for VHS [12].

Deep learning is a technique within the umbrella of artificial intelligence (AI) which is becoming increasingly common in medical imaging to assist radiologists [13, 14]. Convolutional neural networks are a type of deep learning algorithm often used for medical imaging, which learn by analyzing large datasets without the need for extensive programming [13]. Researchers have applied deep learning to several areas within veterinary cardiology [15–17], including convolutional neural network algorithms for detecting cardiomegaly in a binary manner [15, 16], and to measure VHS [18, 19]. A deep learning enabled software from MetronMind (MetronMind, Inc, Paso Robles, California, USA), was developed to measure VHS and VLAS on right lateral thoracic radiographs in dogs. The method used by MetronMind to obtain VHS and VLAS measurements is slightly modified from traditional methods as it uses an average vertebral size which is calculated by taking the mean length of vertebrae four to nine. This modified method has a strong positive correlation with the traditional method described by Buchanan and Büchler [20], although this study only included 42 dogs.

The primary objectives of this study were to compare traditional and modified methods for measuring VHS and VLAS by a trained human observer and to compare to modified measurements by a trained observer with MetronMind for measuring VHS and VLAS on right lateral radiographs from a large diverse population of dogs. We further sought to compare assessment of VHS and VLAS on right and left lateral radiographs made by a trained observer and MetronMind.

## Materials and methods

### MetronMind training set

MetronMind, developed by one of the authors (JJC), is web-based software which uses deep learning to measure VHS and VLAS, it is also available for use within a picture archiving and communication system (PACS) system. The algorithm utilizes convolutional neural network technology to detect objects and measure VHS and VLAS. An initial training set of over 11,000 radiographs annotated by engineers who were trained by veterinarians was used to develop the program. Spot checking was done by veterinarians. Radiographs used in this study were not used for training.

### Dataset

This was a retrospective, single-center comparison study. Medical records of dogs with thoracic radiographs including a right lateral view were retrospectively identified from the Texas A&M Veterinary Medical Teaching Hospital from 2010-2020. Dogs were included if judged by an experienced observer to have a measurable VHS; that is, all landmarks required to measure VHS were identifiable. Where available, left lateral thoracic views were also included. For convenience, three cohorts of dogs who met the criteria above were included in this study: a group of various breeds with myxomatous mitral valve disease (MMVD), a group of Cavalier King Charles Spaniels with and without MMVD [21], and a group of large breed dogs with structurally normal hearts, MMVD, dilated cardiomyopathy and arrhythmogenic cardiomyopathy [22]. Dogs were excluded if the right lateral thoracic radiograph could not be obtained from an eFilm Workstation or if a VHS could not be measured due to vertebral malformations, poor positioning or the inability to visualize all landmarks in a single view. Records were anonymized and missing data were omitted.

### Manual radiograph assessment (human observer)

Lateral radiographs were evaluated using a digital radiography system (eFilm). The quality of six VHS landmarks (central and ventral aspect of the carina, left ventricular cardiac apex, ventral border of the caudal vena cava insertion into the cardiac silhouette, cranial aspect of left ventricle, fourth thoracic vertebrae and the overall thoracic vertebral bodies) and four VLAS landmarks (central and ventral aspect of the carina, dorsal insertion of the caudal vena cava into the cardiac silhouette, fourth thoracic vertebrae and the overall thoracic vertebral bodies) were subjectively assessed on right and left lateral views by a single trained observer based on the degree of difficulty that was required to identify each landmark and classified as excellent, diagnostic or poor (Fig 1). Landmarks were assessed individually as excellent quality when images were of textbook quality, diagnostic when landmarks were easy to identify and judged to allow accurate line placement and poor when landmarks could be identified but were not ideal and may have affected measurement accuracy. Overall VHS and VLAS landmark assessment were classified as excellent if each individual landmark was excellent, diagnostic if landmarks were excellent or diagnostic, and poor if one or more landmark was assessed as poor. Manual VHS and VLAS measurements were determined by a single observer (KTS) after training with a board-certified cardiologist (SGG).

**Fig 1.**
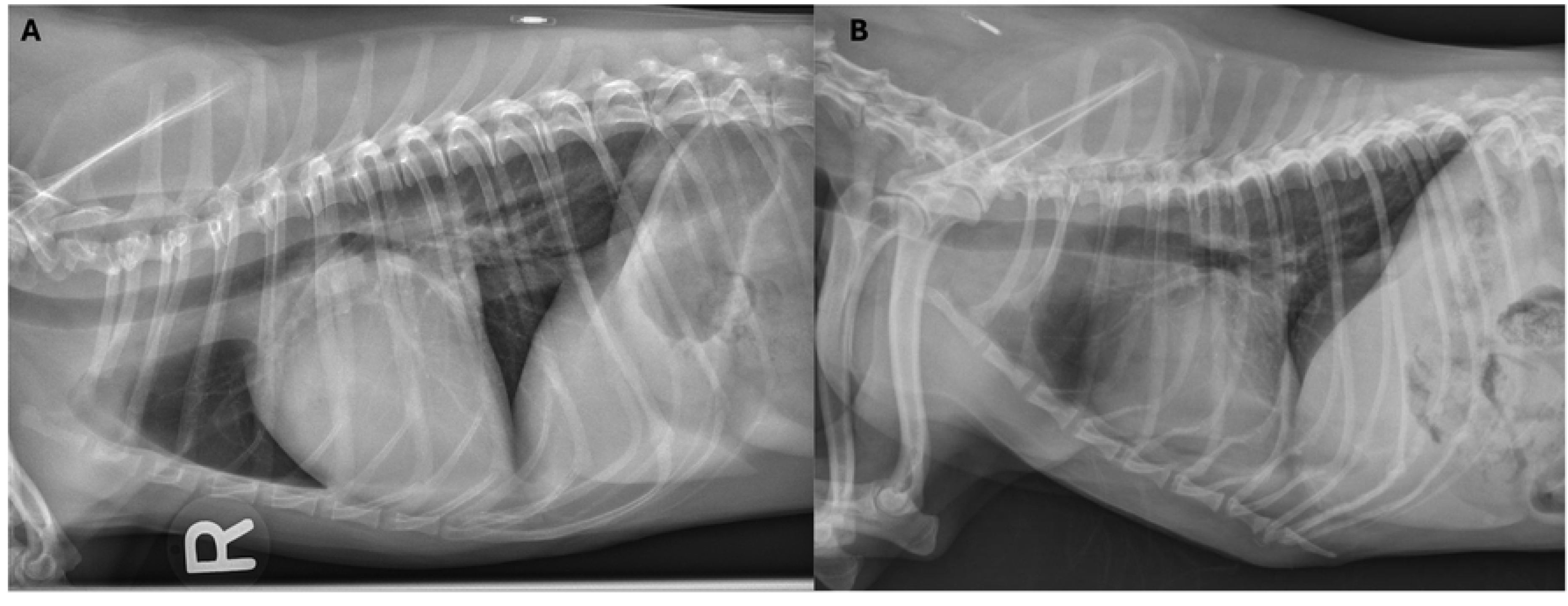
Example of radiographs with variable landmark quality. A) Radiograph with all landmarks classified as excellent. B) Radiograph with poor left ventricular apex and cranial waist; poor vena cava; excellent vertebral bodies and T4; and diagnostic carina.

A traditional VHS (VHS-T) was calculated as follows; the height of the cardiac silhouette was calculated by drawing a line with a digital caliper from the center of the ventral aspect of the carina to the ventral most aspect of the cardiac apex. The carina was identified as the largest circular or ovoid radiolucent structure within the trachea. A second perpendicular line was drawn along the width of the cardiac silhouette starting at ventral aspect of the caudal vena cava and stopping at the cranial waist and perpendicular to the first line. These two lines were transposed along the vertebral column, both starting at the cranial aspect of the 4^th^ thoracic vertebrae. The lengths of both lines were estimated to the nearest 0.1 vertebra (the disc space proceeding a vertebral body is included in the vertebral unit calculation) and then summed (Fig 2A) [5]. Since MetronMind software uses a modified VHS method, whereby an average vertebral length is calculated, a modified VHS method (VHS-M) was also calculated. The cardiac landmarks were kept at the same position. To calculate an average vertebral length a line is drawn across the vertebral bodies, starting at the cranial aspect of the 4^th^ thoracic vertebra and spanning five vertebral bodies to end at the cranial aspect of the 9^th^ thoracic vertebrae. This length (in cm) was divided by five to obtain an average vertebral body length. To calculate the VHS-M, the lengths of the cardiac height and width (in cm) were summed and then divided by the average vertebral body length (Fig 2B).

**Fig 2.**
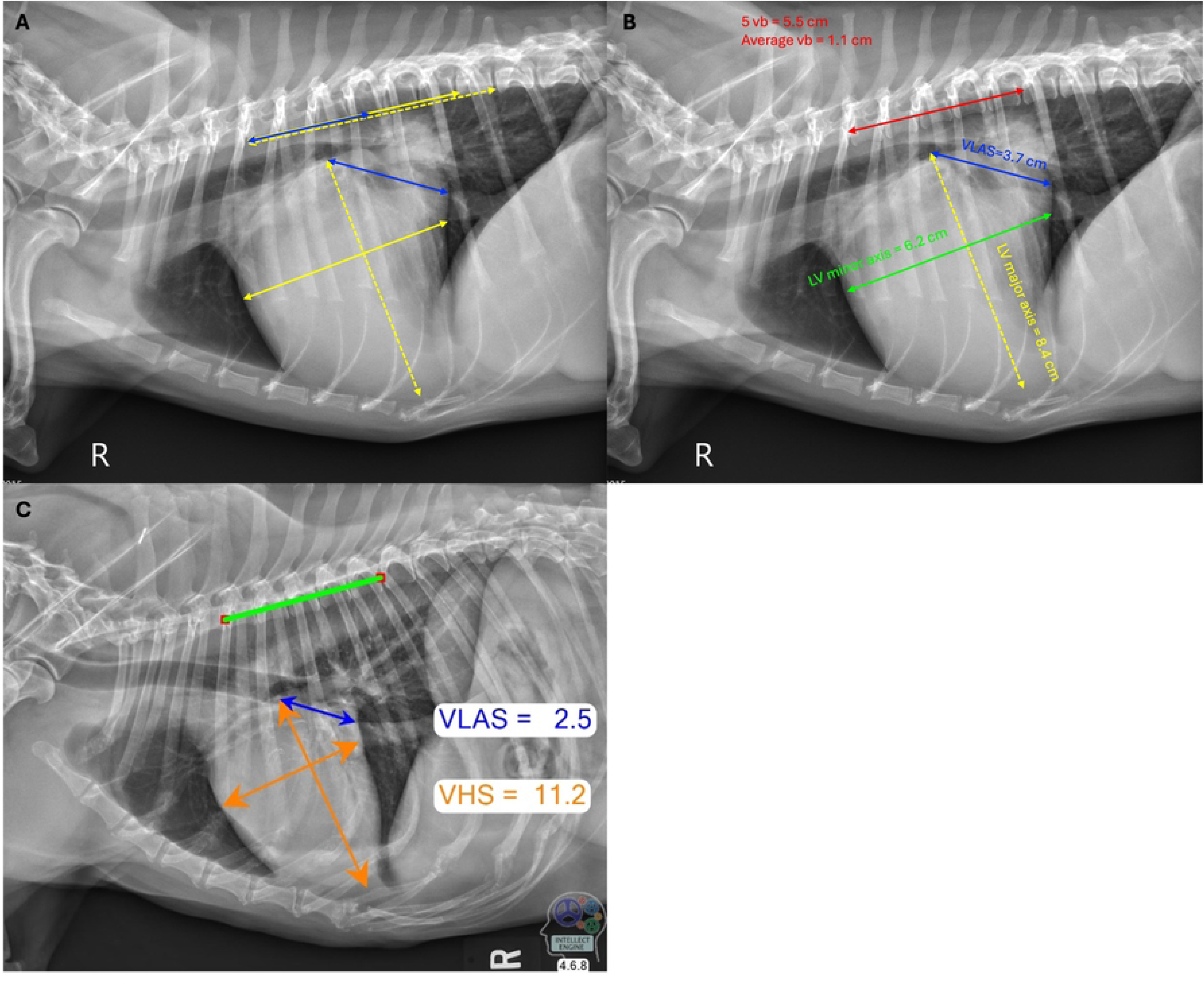
Examples of traditional (A), modified (B) and MetronMind (C) measurements of vertebral heart size (VHS) and vertebral left atrial size (VLAS). A) yellow lines indicate traditional VHS measurements and blue lines indicate traditional VLAS. B) Red line indicates the measurement of five vertebral bodies to calculate modified VHS and VLAS, yellow line the left ventricular (LV) major axis, green line the LV minor axis for VHS and blue line VLAS. C) A green line indicates the measurement of five vertebral bodies, orange lines measures of VHS and blue line VLAS as generated by the MetronMind algorithm.

A traditional VLAS (VLAS-T, Fig 2A) was determined by drawing a line from the center of the most ventral aspect of the carina to the most caudal aspect of the left atrium where it intersects with the dorsal border of the caudal vena cava. Like VHS-T this line was transposed along the vertebral column, starting at the cranial aspect of the 4^th^ thoracic vertebrae, and estimated to the nearest 0.1 vertebra [11]. A modified VLAS (VLAS-M, Fig 2B) was also calculated using the same method of determining average vertebral length as VHS-M.

### Automated radiograph assessment (artificial intelligence algorithm)

Radiographic images were downloaded from the eFilm Workstation and saved as jpeg files. Each radiograph was uploaded into the MetronMind software and analyzed. Once the radiograph was analyzed, an annotated radiograph was produced, displaying the VHS and VLAS landmarks selected by the MetronMind program (version 4.4.3) as well as the deep learning calculated VHS and VLAS values, VHS-AI and VLAS-AI (Fig 2C). The ground truth was defined as the VHS-M and VLAS-M measurements made by a single, trained observer.

### Statistical analysis

Statistical analysis was conducted using R version 4.5.1. Data distribution was examined visually using histograms and using Shapiro Wilk tests. Descriptive statistics for the patient population are reported as median, interquartile range and range. Right lateral radiographic measurements including three VHS and three VLAS values were reported for each enrolled dog: VHS-T, VHS-M, VHS-AI, VLAS-T, VLAS-M and VLAS-AI. Where left lateral radiographs were available, these measurements were also made on the left view (LL-VHS-M, LL-VHS-AI, LL-VLAS-M and LL-VLAS-AI). Correlation between methods was assessed using Spearman correlation. Bland-Altman plots were constructed using the *blandr* package and Passing-Bablok regression using the *mcr* package used to assess agreement between methods. The 95th percentile of absolute difference between VHS-M and VHS-AI were calculated and the 95% confidence interval for this was computed using bootstrapping resampling with replacement using the *boot* package, the number of bootstrap replicates was set to 10,000.

## Results

### Patient population

A total of 1058 dogs representing 80 breeds were included, with the most common being: Cavalier King Charles Spaniel (28.4%), Labrador Retriever (7.9%), Doberman Pinscher (7.8%), and Boxer (6.4%), remaining breeds contributed less than 5% each (S1 Table). There was an equal distribution of males (51.0%) and females (49.0%) and the majority were sterilized (83.8%). Median age was 9.4 years (IQR, 6.7-11.2 years, range, 0.4-17.3 years), and median body weight was 11.6 kg (IQR, 7.5-30.4 kg, range, 1.6-90.0 kg). The range of VHS-T and VLAS-T scores measured by a single trained observer were 8.6-14.7 and 1.2-4.2 demonstrating that a range of cardiac sizes were included in the study. Overall radiographic landmark quality for VHS and VLAS were excellent (15.2% and 21.4%), diagnostic (57.7% and 60.4%) and poor (27.1% and 18.2%, respectively). Quality of individual landmarks was excellent for 46-58% of landmarks while 1.2-13.0% had poor quality (Table 1).

**Table 1:** Classification of individual landmarks. Count and percentage of landmarks classified as excellent, diagnostic, or poor from 1058 radiographs. T4 indicates the fourth thoracic vertebrae.

| Landmark | Excellent | Diagnostic | Poor |
| --- | --- | --- | --- |
| All vertebrae | 518 (49%) | 508 (48%) | 32 (3%) |
| Carina | 610 (57.7%) | 412 (38.9%) | 36 (3.4%) |
| Cranial aspect of T4 | 535 (50.6%) | 510 (48.2%) | 13 (1.2%) |
| Cranial aspect of Left Ventricle | 548 (51.8%) | 457 (43.2%) | 53 (5%) |
| Dorsal vena cava | 486 (45.9%) | 440 (41.6%) | 132 (12.5%) |
| Apex of left ventricle | 487 (46%) | 433 (40.9%) | 138 (13%) |
| Central vena Cava | 544 (51.4%) | 416 (39.3%) | 98 (9.3%) |

### Human: traditional compared to modified

Measurements of VHS-T, VHS-M, VLAS-T and VLAS-M were available from 1057 right lateral radiographic views (S2 table). It was not possible to measure VHS-T on a right lateral radiograph from one case from a giant breed dog as insufficient thoracic vertebrae were present, MetronMind also could not measure VHS-AI. For VHS-M and VHS-T, Pearson’s correlation was 0.994 (95% CI 0.993 to 0.994). Passing-Bablok analysis (Fig 3A) yielded the equation VHS-M=1.06(VHS-T)–0.58, with 95% CI of 1.05-1.07 for the slope and -0.686 to -0.497 for the intercept. Bland–Altman analysis demonstrated a proportional negative bias between VHS-M and VHS-T (Fig 3B), with an overall bias of -0.104 vertebrae (SE 0.004; 95% CI -0.113 to -0.095). For VLAS-M and VLAS-T, Pearson’s correlation was 0.974 (95% CI 0.970 to 0.976). Passing-Bablok analysis (Fig 3C) yielded the equation VLAS-M=1.00(VLAS-T)+0.00, with 95% CI of 1.00-1.00 for the slope and -0.00 to -0.000 for the intercept. Bland–Altman analysis (Fig 3D) showed a bias of 0.040 (SE 0.003; 95% CI 0.033 to 0.046) between VLAS-M and VLAS-T. Comparison between traditional and modified methods for VHS and VLAS were similar for left and right lateral radiographs (S1 Table).

**Fig 3.**
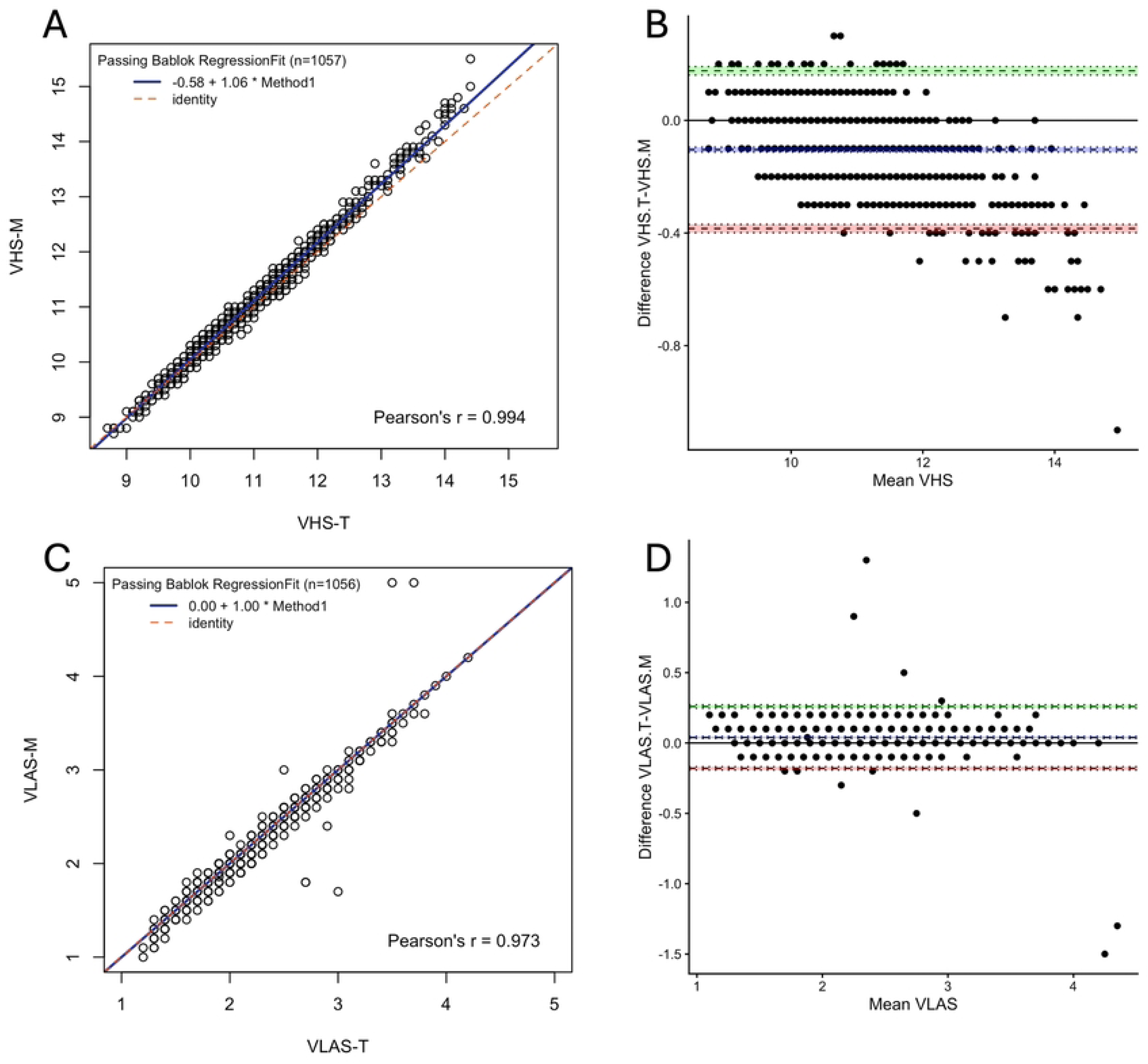
Comparison of traditional and modified methods of measuring vertebral heart size and vertebral left atrial size (VHS-T vs VHS-M and VLAS-T vs VLAS-M) on right lateral radiographs. A) When comparing VHS-T and VHS-M Passing–Bablok analysis yielded the equation VHS-M=1.06(VHS-T)–0.58, Pearson’s linear correlation coefficient r=0.994, n=1055. Blue line: fitted regression line, shading: 95% confidence interval, red dashed line: identity. B) Bland–Altman analysis demonstrated a mean negative bias of −0.10 vertebrae (95% CI −0.11 to −0.10) for the VHS-M method. Dashed lines give limits of agreement (±1.96 SD) and mean bias; shading gives 95% confidence intervals with dotted lines at the limits. C) When comparing VLAS-T and VLAS-M Passing–Bablok analysis yielded the equation VLAS-M=1.00(VLAS-T)–0.00, Pearson’s linear correlation coefficient r=0.974, n=1057. D) Bland–Altman analysis demonstrated a mean bias of 0.040 vertebrae (95% CI 0.032 to 0.046) for the modified VLAS (VLAS-M) method.

### MetronMind compared to human

Measurements of VHS-M, VHS-AI, VLAS-M and VLAS-AI were available from 1057 right lateral radiographic views. For VHS-M and VHS-AI, Pearson’s correlation was 0.947 (95% CI 0.940 to 0.953). Passing-Bablok analysis (Fig 4A) yielded the equation VHS-AI=0.95(VHS-M)+0.44, with 95% CI of 0.95-0.93 for the slope and -0.10 to 0.66 for the intercept. Bland–Altman analysis (Fig 4B) showed a bias of 0.084 vertebrae (SE 0.011; 95% CI 0.062 to 0.104) between VHS-M and VHS-AI. The 95^th^ percentile absolute difference between VHS-M and VHS-AI was 0.70 vertebrae (CI 0.62-0.80, S3 Fig).

**Fig 4.**
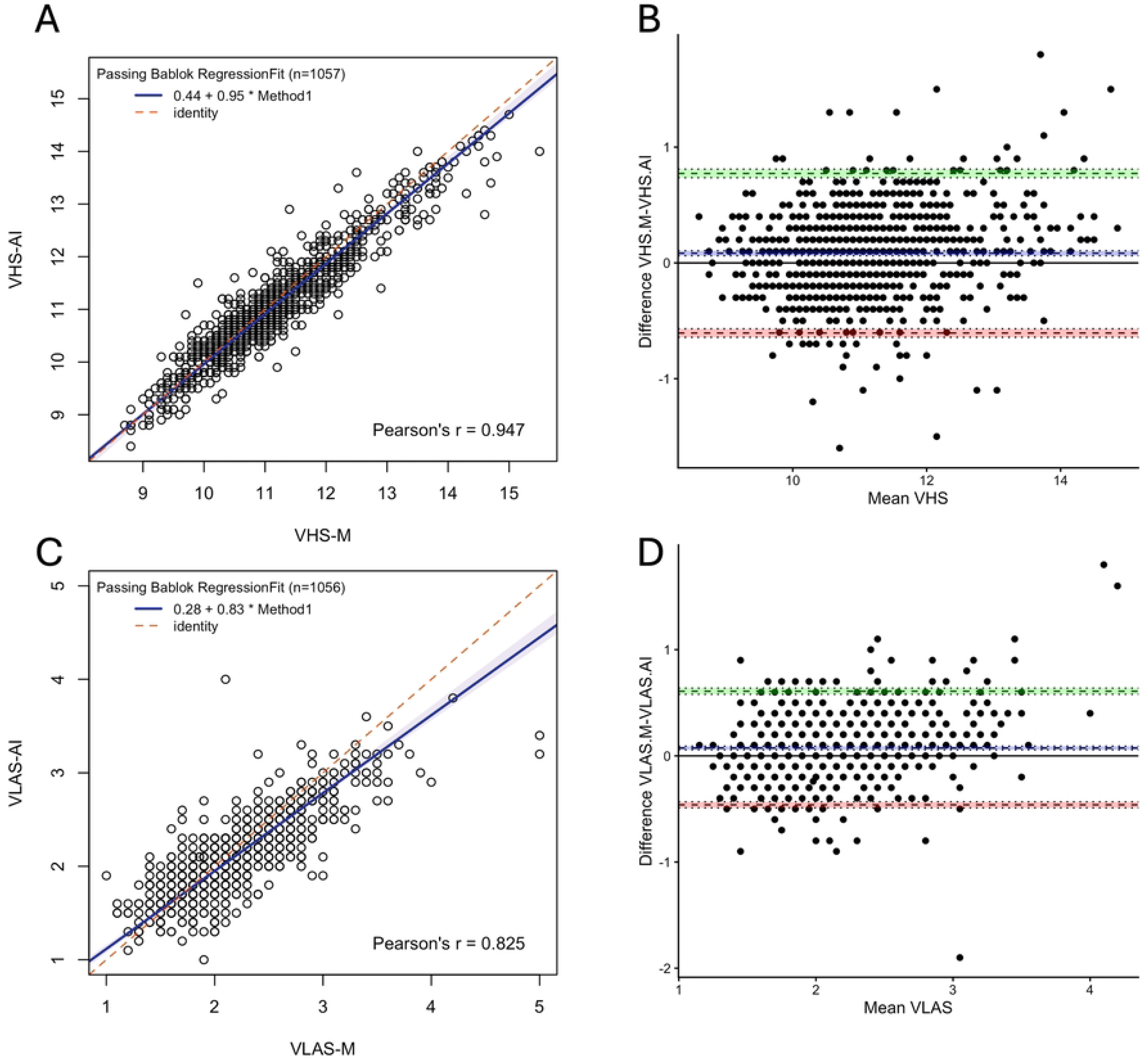
Comparison of modified and artificial intelligence (AI) algorithm (MetronMind) methods of measuring vertebral heart size and vertebral left atrial size (VHS-M vs VHS-AI and VLAS-M vs VLAS-AI) on right lateral radiographs. A) When comparing VHS-M and VHS-AI Passing–Bablok analysis yielded the equation VHS-AI=0.95(VHS-M)+0.44, Pearson’s linear correlation coefficient r=0.947, n=1055. Blue line: fitted regression line, shading: 95% confidence interval, red dashed line: identity. B) Bland–Altman analysis demonstrated a mean bias of 0.08 vertebrae (95% CI 0.06 to 0.10) for the VHS-M compared to the VHS-AI method. Dashed lines give limits of agreement (±1.96 SD) and mean bias; shading gives 95% confidence intervals with dotted lines at the limits. C) When comparing VLAS-M and VLAS-AI Passing–Bablok analysis yielded the equation VLAS-AI=0.083(VLAS-M)+0.28, Pearson’s linear correlation coefficient r=0.811, n=1055. D) Bland–Altman analysis demonstrated a mean bias of 0.07 vertebrae (95% CI 0.05 to 0.09) for VLAS-M compared to the VLAS-AI method.

For VLAS-M and VLAS-AI, Pearson’s correlation was 0.811 (95% CI 0.789 to 0.831). Passing-Bablok analysis (Fig 4C) yielded the equation VLAS-AI=0.83(VLAS-M)+0.28, with 95% CI of 0.80-0.88 for the slope and 0.20 to 0.36 for the intercept. Bland–Altman analysis (Fig 4D) showed a bias of 0.070 vertebrae (SE 0.008; 95% CI 0.053 to 0.087) between VLAS-AI and VLAS-M. Comparisons of modified and AI methods for VHS and VLAS were similar for left and right lateral radiographs (S2 Table). The 95^th^ percentile absolute difference between VLAS-M and VLAS-AI was 0.49 vertebrae (CI 0.44-0.51, S4 Fig).

### Comparison of right and left lateral images

For left and right lateral radiographic views there were 445 measurements of VHS-M and VHS AI. Pearson’s correlation (0.912; CI 0.895 to 0.926) was higher for AI than for humans (0.871; CI 0.847-0.892, Fig 5A), whilst Bland–Altman analysis showed a higher bias (0.31; CI 0.26-0.35) for AI than humans (0.264; CI 0.209 to 0.321, Fig 5B). A total of 439 measurements of VLAS-M and 445 measurements of VLAS-AI were available for left and right lateral radiographic views. Pearson’s correlation was higher for VLAS-M (0.726; CI 0.679 to 0.767, Fig 5C) than for VLAS-AI (0.640; CI 0.582 to 0.692) and Bland–Altman analysis showed a similar negative bias of -0.130 vertebrae (CI -0.164 to -0.096, Fig 5D) for VLAS-M than for VLAS-AI (-0.099; CI -0.131 to -0.067 vertebrae).

**Fig 5.**
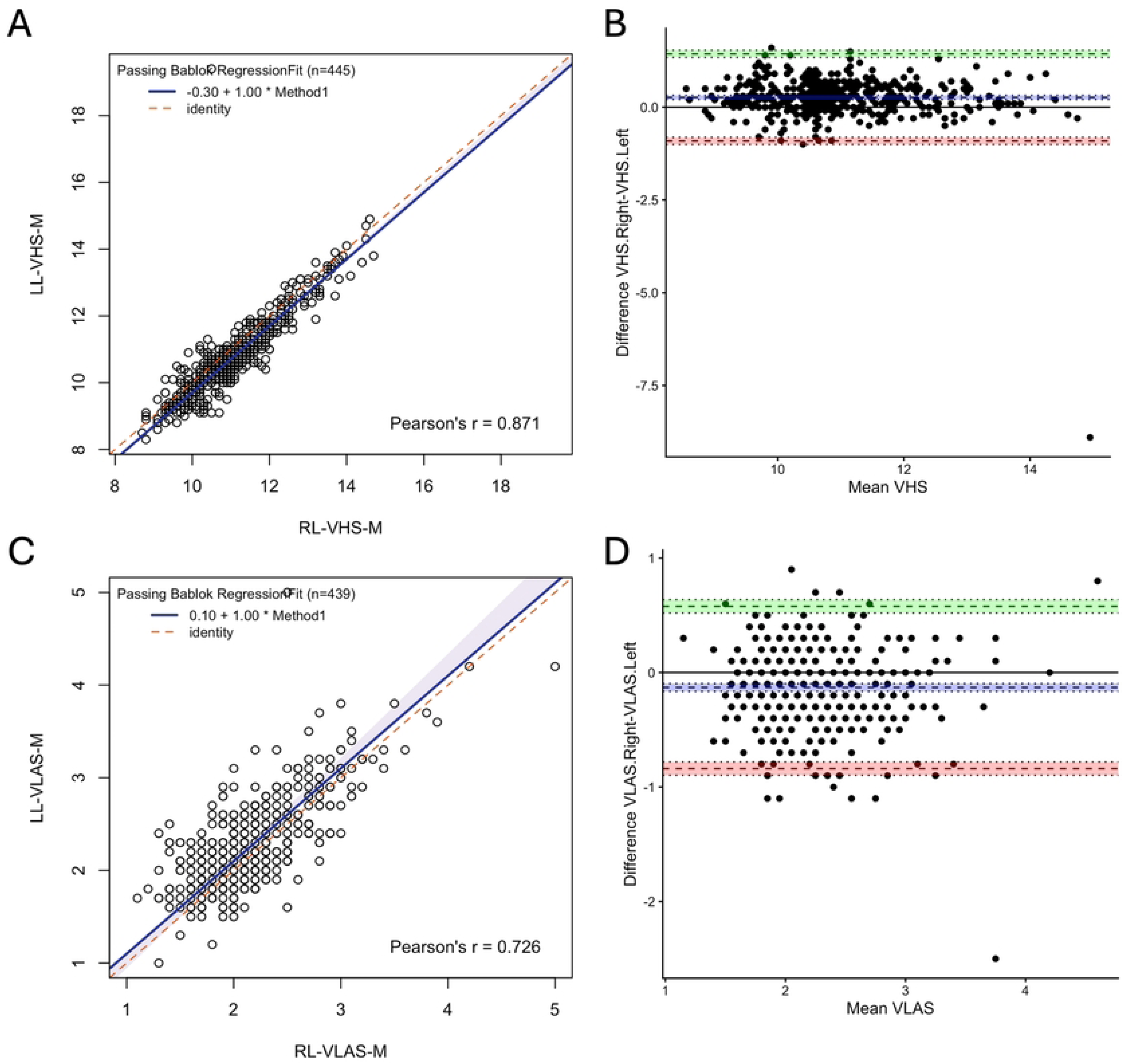
Comparison of measuring modified vertebral heart size (VHS-M) and vertebral left atrial size (VLAS-M) on left (LL) and right (RL) lateral radiographs. A) When comparing measurement of VHS-M on left and right lateral radiographs Passing–Bablok analysis yielded the equation LL-VHS-M=1.0(RL-VHS-M)-0.30, Pearson’s linear correlation coefficient r=0.871, n=445. Blue line: fitted regression line, shading: 95% confidence interval, red dashed line: identity. B) Bland–Altman analysis demonstrated a mean bias of 0.27 vertebrae (95% CI 0.21 to 0.32) for measurements made on the right compared to left radiographs. Dashed lines give limits of agreement (±1.96 SD) and mean bias; shading gives 95% confidence intervals with dotted lines at the limits. C) When VLAS-M on left and right lateral radiographs Passing–Bablok analysis yielded the equation LL-VLAS-M=1.0(RL-VLAS-M)+0.28, Pearson’s linear correlation coefficient r=0.726, n=439. D) Bland–Altman analysis demonstrated a negative mean bias of -0.13 vertebrae (95% CI -0.16 to -0.10) for measurements made on the right compared to left radiographs.

## Discussion

Our study showed minimal differences were found between traditional and modified methods (VHS-T vs VHS-M and VLAS-T vs VLAS-M) that were not of a clinically relevant magnitude. This suggests that cut-offs established using the traditional VHS and VLAS methods should be applicable to the modified method of measurements. The deep learning enabled software, MetronMind, provided VHS and VLAS measurements which are comparable to a trained observer. Finally, VHS and VLAS measurements from right lateral radiographs failed to agree with those from left lateral radiographs sufficiently enough to be interchangeable. Consequently, right lateral radiographs remain the view of choice for measuring VHS and VLAS.

MetronMind uses the average of five vertebral lengths to measure VHS and VLAS, therefore we compared VHS-M and VLAS-M to VHS-T and VLAS-T, which measured the actual number of vertebrae spanned by the cardiac measurements. We found good agreement between VHS-T and VHS-M and between VLAS-T and VLAS-M (mean bias -0.10 and 0.040 respectively), with a strong correlation (0.994 and 0.974 respectively). It is interesting to note that a similar proportional bias was observed for VHS as in another paper [20], whereby higher VHS was associated with a more negative bias. This is likely because VHS-M uses an average vertebral length of T4 to T8, while the traditional method includes thoracic vertebrae beyond T8, which are larger and would result in a lower VHS result. This also explains why we did not observe a proportional bias for VLAS, as all VLAS measurements were below five vertebral units. From a clinical standpoint, various VHS cutoffs have been developed using the traditional VHS method rather than the modified method [5, 23–25]. Given that the results of this study and a previous study [20] demonstrated minimal differences between these methods, the choice of methodology is unlikely to impact interpretation of VHS and previously derived cut-offs can reasonably be applied to a VHS derived by the modified method.

The mean bias derived from Bland-Altman plots between MetronMind and human for VHS and VLAS in this study was 0.08 and 0.07 vertebrae respectively. The small magnitude of this bias means that it is unlikely to cause clinically relevant differences in measurements. The mean bias for VHS is similar to a previous study using AI-algorithms to measure VHS, where measurements by another AI algorithm differed by 0.09 vertebrae compared to board-certified radiologists [18]. Both AI algorithms give less variation than the 1.05 vertebrae that was observed between 16 human observers in another study [6].

Measurements of VHS and VLAS using left and right lateral radiographs had a weaker correlation (0.871 and 0.726 respectively), although still showed a small bias (0.264 and 0.130 respectively). This is consistent with a previous study which demonstrated a higher VHS on right than left lateral projections [26], therefore, the right radiographic projection remains the preferred view.

Echocardiography is considered the most sensitive and specific method for the diagnosis and staging of canine cardiac disease and to measure cardiac size. If echocardiography is unavailable, radiographic indices of heart size, including VHS and VLAS, can be utilized for the diagnosis, staging and monitoring of canine cardiac disease [2, 3, 27]. This study was conducted because echocardiography is not readily available to many practicing veterinarians and there is variability between individuals when measuring VHS and VLAS [6, 12]. A deep learning enabled tool to measure VHS could save time and improve accuracy and consistency in measurements. However, it is important that a human checks placement of radiographic landmarks to ensure diagnostic accuracy. Unlike human observers, deep learning technology has perfect repeatability [6, 12, 28]. Such a tool could therefore be useful for monitoring for cardiomegaly progression. This is particularly useful as the rate of increase in VHS accelerates just before the onset of CHF in MMVD and a rate of change above >0.08 vertebral units per month is associated with the onset of CHF [10, 29]. Similarly, a rate of change of increase in VLAS above 0.02 per month is associated with the onset of CHF in MMVD [30]. MetronMind or other deep learning tools may therefore be particularly useful for repeated measurements and longitudinal monitoring.

A potential concern with utilizing deep learning technology as part of the clinical diagnostic process is the “black box” effect whereby the process of the underlying algorithms remains unclear to the user. Following VHS and VLAS measurements using the MetronMind tool, it is possible to visually assess placement of landmarks and adjust landmarks if needed. While this feature was not used in this study, this could lead to improved performance of the technology in a clinical setting. It is important to note that the algorithm was trained on an initial set of radiographs and performance will be unaffected by use.

The importance of human supervision when deep learning is used in clinical medicine is crucial to emphasize. The algorithm user must be trained to recognize the correct landmarks and review the MetronMind results for accuracy. MetronMind does occasionally make relevant mistakes (such as measuring six vertebral bodies rather than five or severely overestimating the apex). However, with a clinician checking landmark placement, these errors likely would have been caught and corrected. MetronMind can also be deployed within a PACS system on a computer or using an online tool.

The strengths of this study include the use of a single trained observer and a single PACS system for manual measurements, thereby reducing variability that could have been introduced with multiple observers or multiple different measurement systems. This study also included a wide range of heart sizes, body weights and dog breeds, with a variety of thoracic confirmations represented. Additionally, this study included images with landmarks of variable quality, mimicking the realities of clinical veterinary practice. Importantly, when data was analyzed including only radiographs with excellent quality landmarks, or radiographs with excellent and diagnostic quality landmarks, the results were similar to those obtained when all images (including those with poor landmarks) were assessed (S2 Table). This supports the utility of this technology across images of variable landmark quality.

This study also had several limitations. A single observer obtained the manual VHS measurements (VHS-M and VHS-T) in an effort to reduce the impact of interobserver variability [6]; intraobserver variability was not evaluated. This single observer was also considered the ground truth, despite recognition that humans are inherently variable [6, 12]. This study did not include patients with active heart failure or pulmonary infiltrates associated with other etiologies, and pulmonary infiltrates may affect performance. Additionally, some VLAS measurements were unmeasurable due to difficulty in identifying the dorsal aspect of the caudal vena cava landmark. For this study, radiographs were excluded if it was not possible to identify all VHS landmarks, while ability to measure the VLAS was not considered during radiograph selection. This image inclusion criteria could have affected VLAS study results. Lastly, MetronMind will annotate and provide a VHS and VLAS on all lateral projections with the cranial aspect of the thorax displayed to the left and cannot distinguish between right and left lateral views that are displayed in this manner. Similarly, while we cannot predict the performance of MetronMind on non-diagnostic radiographs, the accuracy is likely to be reduced.

Several future studies would be useful to build upon the data presented here. First, factors influencing radiographic measurements like breed, body size or sex were not examined in this study but may warrant specific investigation in the setting of deep learning models. Additionally, investigation into the effect of radiographic pulmonary infiltrates on the performance of MetronMind would be clinically useful for veterinarians assessing dogs with suspected congestive heart failure, where confidence in underlying heart size assessment is particularly relevant for rapid clinical decision making. Future studies comparing the agreement between radiographic measurements from MetronMind with echocardiographic measurements of heart size would also be useful. For example, if echocardiographic Stage B2 MMVD cut-offs were strongly correlated with radiographic assessments from MetronMind, this would be particularly useful and could further support the use of this tool in making clinical decisions for patients with cardiac disease.

In summary, deep learning technology can be used to measure VHS and VLAS using left and right lateral canine thoracic radiographs, although right lateral views remain the preferred view. MetronMind, a commercially available program, is reasonably accurate for measuring VHS and VLAS when compared to a veterinarian and has perfect repeatability [31]. Following AI annotation and measurement, it is possible to manually adjust landmarks if annotations are inaccurate, enabling clinicians to confirm the validity of the measurement and to avoid mistakes in clinical decision making.

## Acknowledgments

This study was presented as an abstract at the 30^th^ ECVIM-CA online congress on September 3, 2020. The authors thank the cardiology faculty, residents and technicians at the Texas A&M University, College of Veterinary Medicine and Biomedical Sciences for their assistance with data collection, and Mark Rishniw and Paul Pion their assistance with initiating the project.

## Supporting information

**S1 Table. Table of dog breeds included in the study.**

**S2 Table. Table of statistical comparisons including Pearson’s correlation, Bland-Altman analysis and Passing-Bablok analysis.**

**S3 Fig. Histogram of the difference between modified vertebral heart size (VHS) determined by human observer (VHS.M) and MetronMind (VHS.AI).** The red lines indicate the 5th percentile and 95th percentile. The blue dashed lines represent the 2.5th percentile and the 97.5th percentile

**S4 Fig. Histogram of the difference between modified vertebral left atrial size (VLAS) determined by human observer (VLAS.M) and MetronMind (VLAS.AI).** The red lines indicate the 5th percentile and 95th percentile. The blue dashed lines represent the 2.5th percentile and the 97.5th percentile

